# Massively Parallel Bead-Free Force Spectroscopy with Fluorescence

**DOI:** 10.1101/2025.09.14.675960

**Authors:** Adam B. Yasunaga, Ryan Riopel, David t. R. Bakker, Dyuti Raghu, Micah Yang, Isaac T. S. Li

**Affiliations:** Department of Chemistry, The University of British Columbia, Kelowna, BC, V1V1V7, Canada

**Author notes:** Authors of equal contribution (Adam B. Yasunaga, Ryan Riopel).

## Abstract

Single-molecule force spectroscopy (SMFS) has transformed our understanding of biomolecular mechanics. However, current high-throughput implementations rely on beads to apply force, introducing size and surface chemistry variability, requiring per-bead calibration, and are prone to multi-tether artifacts. Long handles further complicate measurements by convolving target conformational changes with handle stretching. We introduce Tether Force Spectroscopy (TFS), a bead-free SMFS platform where a single DNA tether acts as both the force applicator and internal calibrator. In TFS, shear flow acting on identical DNA tethers applies piconewton-scale forces directly to surface-anchored molecules whose conformational dynamics are simultaneously monitored by single-molecule fluorescence. This guarantees single-tether results with uniform, internally calibrated forces and is inherently compatible with single-molecule fluorescence. We achieved high-resolution, high-throughput measurements across hundreds of molecules, enabling both force-extension and rupture experiments without specialized instrumentation. The combination of simplicity and simultaneous force-fluorescence capability makes TFS broadly accessible for correlating structure and function in diverse biomolecular systems.

## Introduction

Single-molecule force spectroscopy (SMFS) enables direct manipulation of biomolecules with piconewton-scale forces^1–3^, providing unique insights into protein folding^4–6^, nucleic acid mechanics^7–10^, receptor-ligand binding^11–13^, molecular motor mechanisms^14–16^, and cellular adhesion and mechanotransduction mechanisms^17–20^. These advances highlight the potential of SMFS for uncovering how mechanical forces regulate biological function at the molecular level.

High-resolution SMFS methods such as optical tweezers^21^ and atomic force microscopy^22^ have been central to these studies, but are limited by throughput, as they typically probe one molecule at a time. Bead-based parallelized approaches such as magnetic tweezers^23^, acoustic force spectroscopy^24^, centrifugal force microscopy^25^, and hydrodynamic force spectroscopy^26^ address throughput by enabling simultaneous measurements across many molecules, opening the possibilities of using SMFS as a screening platform to explore diverse nanomechanical behaviours^24–28^. However, several practical challenges continue to limit their adoption. The accuracy of bead-based systems is highly sensitive to intrinsic variability in bead size, requiring per-bead calibration through time-consuming tethered particle tracking in every experiment. The throughputs of these systems are affected by non-specific interactions or multiple tethers that require on-going surface chemistry optimization and extensive post-processing to isolate true single-molecule events^29,30^. Substantial optimization is required for each new system, hindering their overall adoption at a screening scale. Furthermore, the use of long DNA handles are necessary but complicate extension measurements, as bead displacements reflect the convolution of the target molecule’s conformation changes with the tether’s elasticity, obscuring direct observation of the conformational dynamics of the target molecule^31^. Beyond these technical issues, bead-based methods require specialized instrumentation to apply forces on beads, often limiting accessibility and complicating integration with single-molecule fluorescence. Yet, combining single-molecule fluorescence and SMFS provides powerful opportunities to directly correlate force with structural and functional dynamics, as demonstrated in fluorescence optical tweezers studies^32–34^. Incorporating single-molecule fluorescence into massively parallel SMFS platforms remains impractical for most laboratories.

Here, we introduce single-molecule Tether Force Spectroscopy (TFS), a massively parallel bead-free SMFS platform that eliminates these barriers. In TFS, long identical double-stranded DNA (dsDNA) tethers act both as force applicators and internal calibrators under controlled hydrodynamic shear flow, guaranteeing single-tether geometry and uniform force. Critically, TFS inherently uses single-molecule fluorescence for detection, enabling direct observation of conformational changes with nanometer (single-molecule localization) and sub-nanometer (single-molecule FRET) precision. We developed a force calibration model relating shear stress and dsDNA length to the tension at the molecular anchor point. We demonstrated the use of TFS to obtain force-extension curves of DNA hairpin unfolding and quantitatively validated our calibration model by comparing measurements against optical tweezers under identical conditions. Additionally, we used TFS to measure the force-induced dissociation of digoxigenin-antidigoxigenin (Dig-AntiDig) interactions and DNA duplex unzipping. With forces spanning 0-65 pN applied in parallel to hundreds of molecules within a single field of view, TFS combines high resolution, throughput, and fluorescence integration in a single platform, opening new opportunities for high-throughput studies linking molecular mechanics to structure and function. By addressing the key limitations of bead-based SMFS methods, we anticipate TFS will broaden access to SMFS and enable new high-throughput applications across the biomolecular sciences.

## Results and Discussions

### Tether Force Spectroscopy

TFS applies piconewton-scale forces to surface-immobilized biomolecules through hydrodynamic stretching of 8-18 kilobase pair (kbp) dsDNA in a microfluidic channel (Fig. 1a-c). Each DNA tether, anchored at one end, experiences increasing tension along its contour from the free end toward the anchor, with maximum force at the attachment point. The flow-stretched DNA acts as a mechanical transducer, converting shear stress into calibrated tension. Monovalent attachment ensures that each tether exclusively applies force to a single target molecule, enabling controlled and unambiguous force spectroscopy measurement (Fig. 1d).

**Figure 1.**
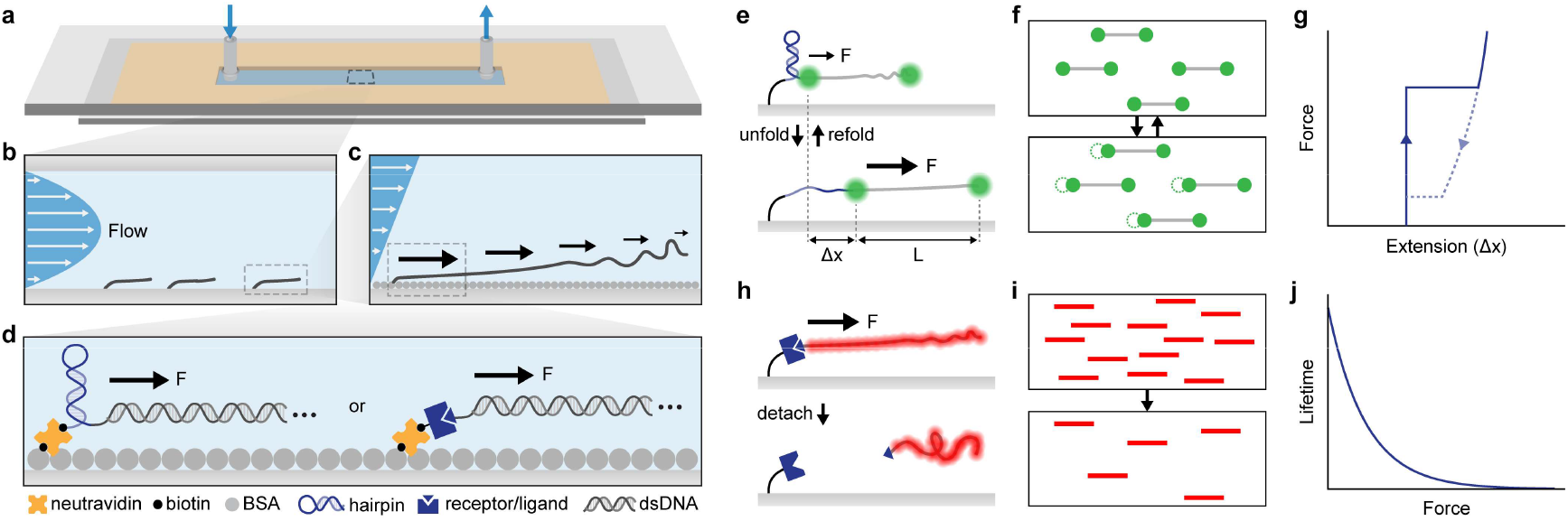
Experiment concept and setup. **a-d,** Schematic of the TFS experimental setup. DNA molecules are immobilized on the bottom surface of a microfluidic channel (**a, b**). Hydrodynamic shear flow exerts tension along each DNA tether, with the greatest tension at the anchor position (**c**). This tension acting at the anchor position can be utilized to exert forces (*F*) on biomolecules (**d**) such as DNA hairpins (left), or receptor-ligand interactions (right). **e, f,** Reversible unfolding experiments. DNA tether length (*L*) and target molecule extension changes (*Δx*) are simultaneously tracked using anchor-end and free-end fluorophores, enabling direct observation of unfolding and refolding transitions. **g,** Schematic of the force-extension curve from DNA hairpin unfolding measurements showing molecular extension due to the hairpin only. **h, i,** Irreversible rupture experiments, in which rupture events result in detachment of DNA from the surface. **j,** Schematic force-dependent lifetime resulting from irreversible rupture experiments. DNA lengths are not to scale.

To study both reversible conformational changes and irreversible rupture events, we established two complementary experimental modes. For reversible structural transitions, such as DNA hairpin unfolding, precise detection of nanometer-scale extension changes is achieved by labelling DNA at both anchor and free ends with single fluorophores (Cy3 or Cy3B), directly imaged by Total Internal Reflection Fluorescence (TIRF) microscopy (Fig. 1e). Single-molecule localization of these fluorophores simultaneously reports the anchor displacement (*Δx*) and total tether length (*L*) (Fig. 1e). Unfolding and refolding transitions can be directly identified from *Δx*, which can be correlated with calibrated tether forces derived from *L* to generate precise force-extension curves for each molecule (Fig. 1f, g). Unlike bead-based methods, all TFS tethers are identical dsDNA molecules, ensuring uniform force application and eliminating uncertainty from force applicator heterogeneity.

For irreversible events, such as receptor-ligand dissociation, ruptures are detected by the loss of fluorescent DNA tethers. DNA tethers labelled with intercalating dye (e.g., SYTOX Deep Red) appear as lines tethered to the surface (Fig. 1h, i). Upon rupture of the anchoring interaction, the tether disappears as it is swept downstream by flow. Using a shear stress-to-force calibration, the shear stress at rupture provides the rupture forces. This binary readout enables simple tracking of hundreds of molecules simultaneously, allowing efficient determination of force-dependent lifetimes (Fig. 1j).

Together, these two modes establish TFS as a platform for both reversible and irreversible SMFS assays, combining high-resolution, uniform and internally calibrated force, and high-throughput, while avoiding limitations of bead-based systems.

### Force Calibration of DNA Flow-Stretching

Accurate SMFS requires a robust calibration linking the experimentally controlled input (shear stress) to the force applied to the target molecule. In TFS, hydrodynamic flow in the microfluidic channel creates shear stress, resulting in drag forces distributed along the DNA tether. The combination of shear stress and tether length determines the total force at the anchor.

To establish this relationship, we tracked fluorophores at both the anchor and free ends of individual DNA molecules under a series of shear stresses (Fig. 2a-f, Fig. S1a). When imaged with TIRF microscopy, the anchor fluorophores remained stationary, while the free-end fluorophores shifted along the flow direction with increasing shear stress (Fig. 2b). At higher shear stress, greater force and extension reduced free-end motion, resulting in brighter fluorescence signals (Fig. 2b). Single-molecule localization of both fluorophores across shear stresses from 0.4 to 11 Pa provided more quantification of this behaviour (Fig. 2c-f). The anchor fluorophore formed a tight, shear-independent localization cluster (radial S.D. = 8.5 ± 2.6 nm, Fig. 2d, e; Fig. S2a, b). In contrast, the free-end fluorophore shifted progressively in the flow direction and exhibited reduced positional spread, from ∼80 to ∼6 nm at the low-end and high-end of shear stresses (Fig. 2d, e; Fig. S2c, d). A super-resolved image combining localization data across all shear stresses further illustrates the distinct behavior of the anchor and free-end fluorophores (Fig. 2f, Fig. S2e, f).

**Figure 2.**
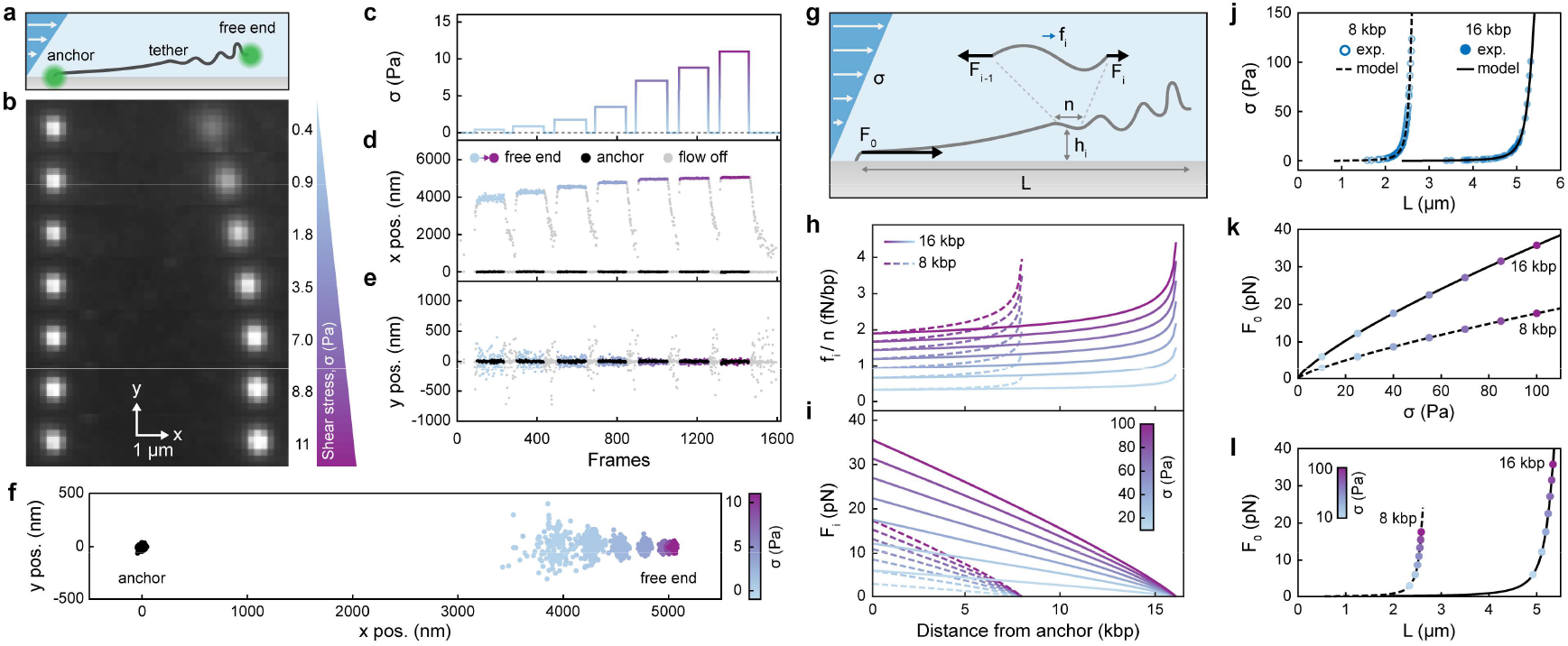
DNA flow-extension measurements and model. **a,** Schematic of DNA length measurement under flow, with a 16 kbp DNA tether terminally labelled with Cy3 (anchor) and Cy3B (free end). **b,** TIRF images of 16 kbp DNA stretching at various shear stresses (values in Pa shown at right). **c,** Shear stress profile applied over the course of an experiment. **d, e,** x-(**d**) and y-(**e**) positions of localizations from the molecule shown in **b**, in response to the applied shear stresses in **c. f,** Super-resolved image of the molecule in **b** shown across the range of applied shear stresses, with x*-* and y-positions corresponding to those plotted in **d** and **e. g,** Schematic of the segment-based model, where the drag force on each DNA segment (*f_i_*) is calculated from shear stress (*σ*), segment length (*n*), and segment height (*h_i_*); cumulative tension (*f_i_* or *F_0_* at the anchor) is computed by summing segmental forces. **h,_i_,** Model-predicted profiles of drag force (**h**) and cumulative tension (**i**) along the contour of 8 kbp and 16 kbp DNA under various shear stresses. **j,** Experimentally measured DNA length as a function of shear stress overlaid with model predictions for both 8 kbp and 16 kbp constructs. **k, l,** Calibration curves extracted from the model, showing *F_0_* as a function of applied shear stress (**k**) or measured DNA length (**l**).

Because tension along a flow-stretched DNA is non-uniform, increasing from the free end to the anchor^35,36^, conventional uniform-tension models such as the extensible worm-like chain (XWLC) cannot be applied directly. Instead, we modelled DNA as a series of discrete segments, each experiencing a drag force *f_i_* and downstream tension *F_i_*, balanced by upstream tension *Fi-1* (Fig. 2g-i). This produces a continuous tension gradient from zero at the free end to a maximum force at the anchor (Fig. 2i). With shear stress as the input and tether length as the readout (Fig.2j), the model provides an empirical calibration of anchor force (*F_0_*) as a function of either shear stress (*σ*, Fig. 2k) or total tether length (*L*, Fig. 2l). A single set of parameters accurately fit both 8 and 16 kbp DNA tethers (Fig. 2j), underscoring the model’s generalizability. These two complementary calibrations enable precise control over force application through shear stress, and per-molecule force determination from tether length, as demonstrated in the following sections.

### DNA Hairpin Unfolding with Integrated Force and Fluorescence

With the force calibration established, we next applied TFS to measure force-induced unfolding and refolding of DNA hairpins. A 16 kbp DNA tether terminally labelled with Cy3B was anchored to the surface via a biotinylated DNA hairpin consisting of a 15 bp, 60% GC stem and a 30 nt poly-T loop (Fig. 3a; Fig. S1b). Force ramps were generated by applying a triangular waveform of shear stress (Fig. 3b), driving repeated unfolding and refolding cycles.

**Figure 3.**
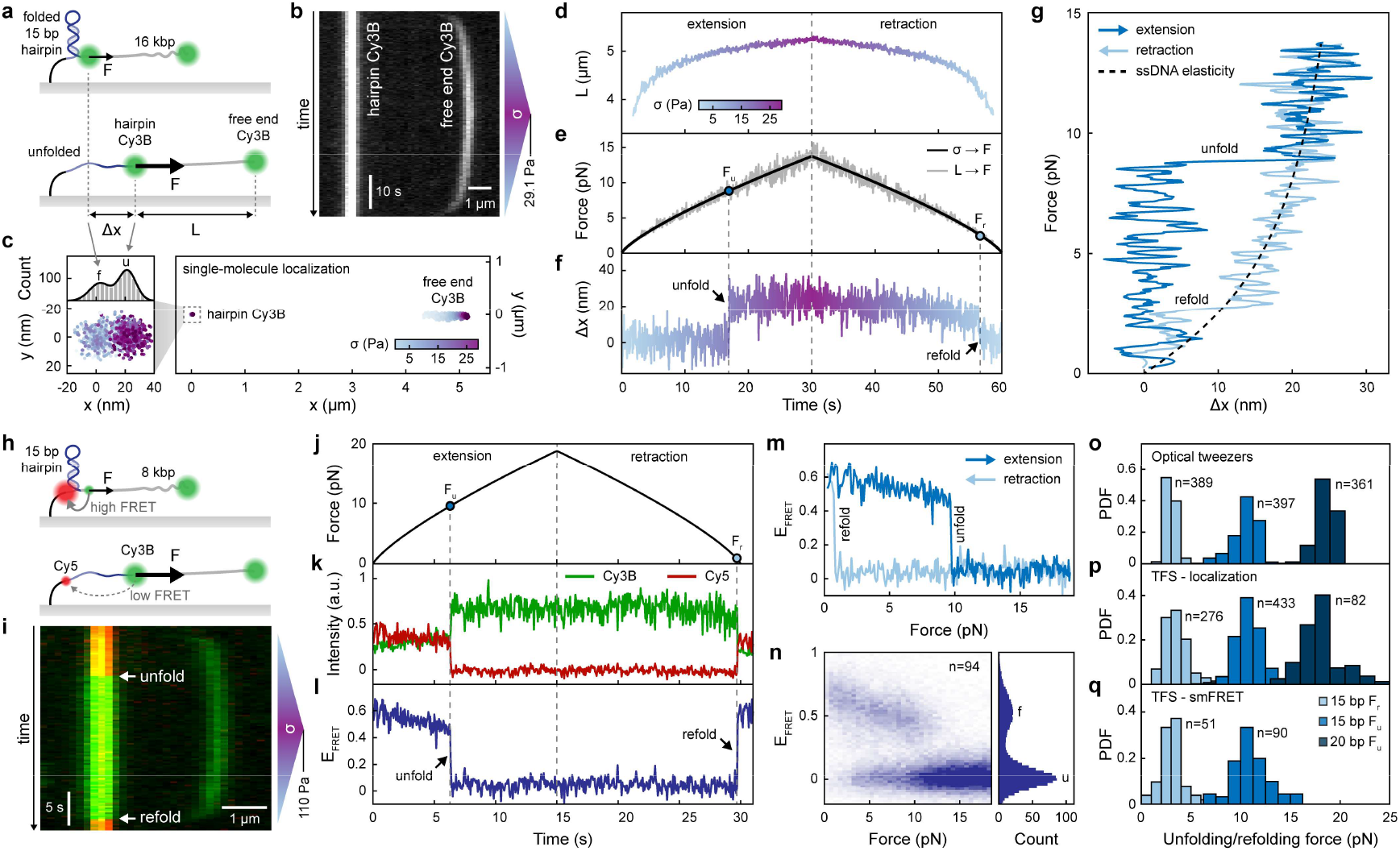
DNA Hairpin Unfolding Measurements. **a,** Schematic of DNA hairpin unfolding experiments, in which a force (*F*) is applied to a hairpin through terminally labelled 16 kbp DNA, where unfolding is measured through hairpin extension (*Δx*) as 16 kbp DNA length (*L*) is monitored. **b, c,** Kymograph (**b**) and resulting localization (**c**) of 16 kbp DNA stretching, colour-coded by shear stress, with a bimodal distribution of localizations corresponding to the unfolded and folded states. **d, e, f,** Data from the molecule in **c** plotted against time, colour-coded by shear stress, exhibiting symmetrical changes in length (**d**), which can be utilized to determine force over time (**e**, shown in grey), in addition to force measurements determined from shear stress (**e**, shown in black). Unfolding and refolding forces (*F_u_* and *F_R_*) are identified from discontinuities in hairpin fluorophore x-position (**f**). **g,** Force-extension curve produced from the data in **e** and **f** (with smoothing applied), overlaid with the stretching of the 60 nt ssDNA released by unfolding as predicted by the XWLC model. **h,** Schematic of smFRET hairpin unfolding experiments, in which terminally labelled 8 kbp DNA is anchored through a hairpin labelled on opposite ends of its stem with Cy5 and Cy3B fluorophores. **i,** Multi-channel kymograph of 8 kbp DNA stretching, exhibiting anti-correlated changes in Cy5 (red) and Cy3B (green) intensities. **j, k, l,** Example smFRET data from the molecule in **i**, demonstrating changes in donor and acceptor intensities (**k**) in response to the applied force (**j**), which can be utilized to determine FRET efficiency (*E_FRET_*, **l**). **m, n,** FRET efficiency as a function of force for a single molecule (**m**), and an ensemble of molecules (**n**). **o, p, q,** Distributions of measured unfolding and refolding forces acquired through dual-trap optical tweezers (**o**), localization-based TFS measurements (**p**), and smFRET-based TFS measurements (**q**). See Fig. S6 for the distribution of applied loading rates. All measured distributions across methods are statistically equivalent within a threshold of ±1 pN as determined by equivalence testing (*p* < 0.05). DNA lengths are not to scale.

Conformational changes were detected by single-molecule localization of the anchor fluorophore adjacent to the hairpin (Fig. 3c). Unfolding caused the fluorophore to displace suddenly in the direction of flow, while refolding reversed this displacement. Unlike the unimodal anchor distributions in Fig. 2f, super-resolved images during force ramps revealed two distinct populations along the x-axis, separated by ∼20 nm, consistent with the expected extension change for hairpin unfolding. Time trajectory of the x position shows discrete, single-step transitions upon both force increase and decrease (Fig. 3f), characteristic of two-state hairpin unfolding and refolding^7^.

Simultaneous tracking of the free-end fluorophore provides DNA tether length (*L*, Fig. 3d) and per-molecule force using the length-to-force calibration (Fig. 2l, 3e). These force values closely match those predicted from the shear stress-based calibration (Fig. 2k, 3e), confirming consistency with model predictions. Plotting force (Fig. 3e) against anchor displacement (Fig. 3f) yields the force-extension curve (Fig. 3g), where the unfolding (∼9 pN) and refolding (∼3 pN) events are evident from abrupt extension changes. Furthermore, the stretching of the unfolded state follows the expected entropic elastic behaviour of the 60 nt ssDNA released upon hairpin unfolding^37^, validating the observed transition.

Our TFS setup achieves ∼8 nm resolution for detecting force-induced conformational changes by single-molecule localization (Fig. S4). To extend resolution, we integrated TFS with single-molecule FRET (smFRET), enabling sub-nanometer detection of structural transitions. An 8 kbp DNA tether containing the same hairpin was modified with a Cy5-labelled oligonucleotide hybridized to the ssDNA between the hairpin and surface, placing Cy5 and Cy3B on opposite ends of the hairpin stem (Fig. 3h; Fig. S1b). FRET efficiency changes reported hairpin state (Fig. 3i-l), with unfolding and refolding transitions consistent with localization data. Notably, smFRET measurements resolved a gradual decrease in FRET efficiency prior to unfolding, suggesting an elastic stretching of the DNA between Cy3B and Cy5 in the folded hairpin state (Fig. 3k-n). This demonstrates the potential to use TFS’s single-molecule fluorescence capabilities to detect subtle nanomechanical behaviours that may be otherwise obscured in conventional methods

Beyond demonstrating the utility of TFS as an SMFS method, these hairpin unfolding experiments were used to validate TFS force calibrations. Unfolding and refolding force distributions measured by optical tweezers, TFS with single-molecule localization-based force readouts, and TFS with smFRET, agreed within experimental precision (Fig. 3o-q; Fig. S5).

Importantly, this agreement held across DNA tethers of different lengths (16 kbp for localization TFS and 8 kbp for smFRET TFS), highlighting the accuracy and robustness of our force calibration. Loading-rate dependence of unfolding and refolding forces was consistent between optical tweezers and TFS (Fig. S6), and direct comparison of extension changes also showed close agreement (Fig. S7). Together, these results confirm the robustness of TFS-based SMFS.

### High-Throughput Quantification of Rupture Kinetics

To demonstrate the ability of TFS to quantify force-dependent dissociation kinetics, we performed irreversible rupture experiments on two systems: Dig-AntiDig interactions and DNA duplex unzipping (Fig. 4a, b; Fig. S1c, d). For Dig-AntiDig rupture measurements, 17.6 kbp DNA was tethered to an AntiDig-coated surface via a 5′ digoxigenin modification (Fig. 4a). For DNA unzipping, 12 kbp DNA was hybridized in an unzipping geometry to an 18 bp 83% GC surface-anchored strand (Fig. 4b).

**Figure 4.**
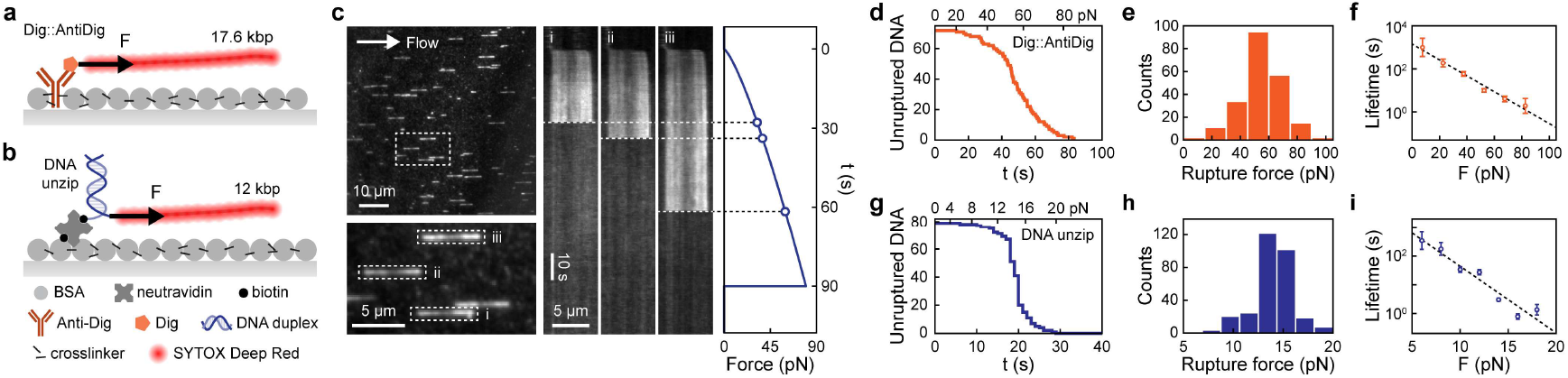
Irreversible Rupture Measurements. **a, b,** Schematic of Dig-AntiDig rupture (**a**) and DNA unzipping (**b**) TFS experiments. **c,** Example frame of Dig-AntiDig anchored 17.6 kbp DNA (left). Corresponding kymographs are shown (center), in which the disappearance of fluorescence can be related to a corresponding rupture force (right). **d, e, f,** Dig-AntiDig rupture data, acquired at a mean instantaneous loading rate of 0.9 pN/s (S.D. 0.1 pN/s), showing the decay in the unruptured population as force is increased over time over a single force ramp (**d**), the rupture force distribution (**e**), and the resulting force-dependent lifetime (**f**). **g, h, i,** DNA unzipping data, acquired at a mean instantaneous loading rate of 0.6 pN/s (S.D. 0.2 pN/s), showing the decay in the unruptured population as force is increased over time over a single force ramp (**g**), the rupture force distribution (**h**), and the resulting force-dependent lifetime (**i**). Error bars on **f** and **i** are standard deviations, computed as described by Zhang and Dudko39. DNA lengths in **a** and **b** are not to scale.

During force ramps, DNA was visualized using SYTOX Deep Red (Fig. 4c). Non-uniform intensity along the DNA reflects preferential intercalation at AT-rich regions (see Fig. S8). Rupture events were identified by the sudden loss of fluorescence intensity along the entire DNA length, and the corresponding rupture force was extracted from the timepoint of rupture along the applied force ramp (Fig. 4c). As the force increases, the number of molecules within the field of view decays to zero within a single experiment, providing statistics from ∼ 100 rupture events in parallel (Fig. 4d, g). The resulting rupture force distributions (Fig. 4e, h) are consistent with previously reported values for these interactions (Supplementary Note 1)^24,26,38^.

These rupture forces may be further utilized to extract physically meaningful parameters such as force-dependent lifetimes that characterize the mechanical strengths of interactions. Using a transformation derived by Zhang and Dudko^39^, rupture force distributions were converted into lifetimes spanning the detected force range (Fig. 4f, i). These lifetimes fit well to exponential decays predicted by the Bell-Evans model^40,41^, confirming the quantitative reliability of TFS for non-equilibrium kinetic measurements.

### A Bead-Free Paradigm for SMFS

Advances in SMFS have traditionally focused on optimizing one capability at a time – combining fluorescence detection, increasing throughput, or improving accessibility. TFS achieves all three goals simultaneously, providing high-resolution (Supplementary Note 2), high-throughput, and single-tether measurement by design. These gains arise from a conceptual shift away from bead-and cantilever-based platforms. Whereas conventional methods apply force and report displacement through micron-sized objects tethered to the molecule of interest, TFS removes this intermediary. The DNA tether itself acts as both the force transducer under hydrodynamic flow and a nanoscale reporter observable by single-molecule fluorescence. This direct actuation and observation scheme removes bead heterogeneity and decouples measurements from handle mechanics.

Single-molecule fluorescence in TFS enables nanometer precision tracking of any labelled sites on the target molecule. Therefore, TFS not only can generate force-extension curves that completely decouple from long handle elasticity (Fig. 3g), but also enable unconventional force-conformation curves that probes structural changes inside of the target molecule (Fig. 3m). Beyond this, the native fluorescence compatibility permits measurements that are impractical with bead-based systems, such as multiplexing force readout along the tether and off-axis conformational tracking.

Flow-stretching DNA further addresses limitations inherent to bead-based force generation. Identical DNA tethers yield uniform forces across target molecules, while tether length itself provides an intrinsic calibration for each molecule. The direct shear stress-to-force relationship enables passive force-clamp experiments with a simple programmable syringe pump. The useful dynamic range of TFS spans up to the overstretching limit of dsDNA (∼65 pN, Fig. S9)^42^, as with all other SMFS methods that utilize DNA handles. At the lower force range, TFS is not constrained by bead thermal motion or trap stiffness. Modelling indicates that in principle, femtonewton-level forces are achievable using short tethers and low shear stress (e.g., ∼10 fN for a 1.5 kbp tether at 50 mPa shear stress).

The two experimental modes of TFS highlight complementary strengths. Force-extension assays require precise single-molecule tracking and demonstrate the resolution of the method, while rupture assays leverage the binary disappearance of stained DNA for scalability and throughput. In the latter, high-intensity intercalator signals relax detection requirements, enabling massively parallel assays with even epifluorescence illumination and minimal post-processing. And most importantly, guaranteed single-tether geometry removes the need for extensive filtering, a major bottleneck in bead-based methods.

Finally, accessibility remains a barrier for high-throughput SMFS adoption. Although methods such as acoustic force spectroscopy and centrifugal force microscopy reduce instrument cost, they still require specialized equipment or modification of existing equipment during implementation, presenting a major hurdle to non-expert labs. In contrast, TFS runs on standard TIRF or epifluorescence microscopes (depending on the mode) with syringe pumps and flow chambers that are common in many research labs, with no modification required. Its simplicity and optical readout make TFS readily compatible with a wide variety of fluorescence-based techniques, enabling studies of biomolecular systems under force with an array of different multiplexing capabilities and detection schemes.

## Conclusion

In summary, TFS addresses long-standing bottlenecks in SMFS by eliminating beads and cantilevers, ensuring single-tether geometry, and directly integrating with single-molecule fluorescence for high-throughput, high-resolution detection of molecular conformations and binding states. It delivers the precision and control valued by specialists with throughput and accessibility needed for wide adoption. By lowering the technical barrier while enabling multimodal readouts, TFS is a versatile platform for probing mechanosensitive biomolecules and has the potential to broaden the reach of quantitative force spectroscopy across the molecular biosciences.

## Methods

### TFS DNA Construction

All DNA oligonucleotides were ordered from Integrated DNA Technologies (Coralville, IA, USA). Sequences are shown in Supplementary Table 1.

8 kbp (8066 bp), 12 kbp (12330 bp), 16 kbp (16173 bp), and 17.6 kbp (17587 bp) PCR products were amplified from a Lambda genome template using LongAmp Taq DNA polymerase (New England Biolabs, Whitby, ON, Canada). PCR products were gel extracted to remove non-specific amplicons and ensure DNA lengths were uniform using the Monarch DNA Gel Extraction Kit (New England Biolabs, Whitby, ON, Canada).

Functionalization of DNA constructs was achieved through modifications to PCR primers, those being a 5’ biotinylated ssDNA oligonucleotide connected through a triethylene glycol spacer for use in calibration and DNA unzipping experiments, a BsaI restriction site for use in hairpin unfolding experiments, and a 5’ digoxigenin for use in Dig-AntiDig rupture experiments. For fluorescent labelling, primers were modified with single Cy3 fluorophores or amine groups that were conjugated to Cy3B NHS ester (Lumiprobe, Westminister, MD, USA). Hairpin constructs were further modified through digestion with BsaI at 37 °C for 1 hour to generate a 4 nt sticky end, which was purified using a Monarch PCR and DNA Cleanup Kit (New England Biolabs, Whitby, ON, Canada), and subsequently ligated to a hairpin oligonucleotide with 5’ phosphate and 3’ biotin modifications at a 1:100 molar ratio for 30 seconds using an Instant Sticky-End Ligase Master Mix (New England Biolabs, Whitby, ON, Canada). Oligonucleotides and assembled DNA constructs were stored at -20 °C.

### Microfluidic Channel Construction

25 × 75 × 1 mm VWR microscope slides were engraved with a BOSS LS-1416 laser engraver to form holes that would later serve as inlets and outlets for each channel. These laser-engraved slides were sonicated along with No. 1.5 VWR coverslips in a saturated solution of KOH in ethanol for 3 minutes, rinsed with methanol and water, dried with N2 gas, and treated with O2 plasma for ∼1 minute in a Harrick PDC-32G plasma cleaner.

Channels were then assembled by sandwiching F9460PC 3M Adhesive Transfer Tape with custom 400 μm wide channels made with a Cricut Explore Air 2 cutter in between the cleaned slide and coverslip, ensuring the laser-engraved holes aligned with the channel ends. Inlet and outlet tubes were attached by connecting E-3606 Tygon tubing to the engraved holes through cut P20 pipette tips, and sealing the connection with Raidzap thick UV resin. Each assembled chip contains 4 separate channels for parallel sample preparation, with each channel being approximately 8 mm long, 400 μm wide, and 60 μm tall.

### Surface Treatment

Prior to the addition of liquid, Ar plasma was injected into channels using a Plasma Pipette (Femto Science, Hwaseong, Republic of Korea) on “High” for ∼10 s in each channel inlet. This was followed by sequential incubations with 5 M aqueous KOH for 15 minutes, 1 mg/ml BSA (Tocris Bioscience, Bristol, UK) and 0.2 mg/ml BSA-biotin (Thermo Scientific, Waltham, MA USA) for 30 minutes, 0.1% glutaraldehyde crosslinker (Sigma Aldrich, St Louis, MO, USA) for 30 minutes, 0.2 mg/ml neutravidin (ThermoFisher Scientific, Waltham, MA USA) for 15 minutes, and 25 - 200 pM DNA tethers for 30 minutes (depending on the desired surface density), with wash steps using water before every incubation except the addition of glutaraldehyde. For calibration and DNA unzipping experiments, a 10 nM solution of the 5’ biotinylated ssDNA oligonucleotide was incubated in the channel for 30 minutes before the addition of DNA tethers (DNA tethers were hybridized in a shearing configuration for calibration experiments, and an unzipping configuration for unzipping experiments). For Dig-AntiDig rupture experiments, the surface treatment protocol was modified in the following respects: an additional 1-hour incubation with 20 μg/ml polyclonal AntiDig (Roche Diagnostics GmbH, Mannheim, Germany) was performed following the KOH incubation, BSA-biotin was removed from the 1 mg/ml BSA incubation, and the neutravidin incubation was not included.

### TFS Experiments

All TFS experiments were performed at room temperature (20 °C). Imaging buffers contained a GODCAT oxygen scavenging system (0.4% D-glucose, 1.12 mg/ml glucose oxidase, 1 mg/ml catalase; Sigma Aldrich, St Louis, MO, USA), 2 mM Trolox triplet-state quencher (Sigma Aldrich, St Louis, MO, USA), and 50 mM tris, with salt concentrations of 200 mM NaCl in the hairpin and Dig-AntiDig experiments, and 150 mM NaCl and 5 mM MgCl2 in the calibration and DNA unzipping experiments. SYTOX Deep Red nucleic acid stain (Invitrogen by Thermo Fisher Scientific, Eugene, OR, USA) was used in imaging buffers for irreversible rupture experiments at 50 nM (Dig-AntiDig experiment) or 1-10 nM (DNA unzipping experiments) to minimize intercalation into the 18 bp unzipping duplex.

Single-molecule TIRF imaging was performed on an Olympus IX83 inverted microscope with an Andor iXon Ultra 897 EMCCD camera. Channel width was determined from 4× brightfield images. Channel height was measured as the axial displacement of the objective between bottom and top channel surfaces in focus and multiplied by an empirically determined factor of 0.8 to correct for refractive index mismatch^43,44^.

For experiments, the bottom surface near the channel center was imaged at 100× in TIRF mode while buffer flow was applied via a Pump 11 Elite Harvard syringe pump controlled by a custom LabVIEW interface. Buffers were delivered from a 5 ml glass Hamilton syringe through BD polyethylene tubing. Cy3/Cy3B-labelled DNA was excited with a 532 nm laser (Spectra-Physics, Excelsior 532), and SYTOX Deep Red-stained DNA was excited with a 642 nm laser (Spectra-Physics, Excelsior One 642).

### Image Processing and Analysis

Movies from calibration and hairpin unfolding experiments were processed using the Picasso software package^45^ to generate single-molecule localizations of each fluorophore in every frame. These localizations were acquired by fitting to the intensity profile of each molecule to determine the maximum likelihood estimate of the x- and y-positions. For hairpin unfolding data, drift correction was performed using stationary debris spots as fiducial markers. Fiducials were manually selected and verified to avoid misidentifying flow-stretching molecules as drift references. After drift correction, anchor and free-end fluorophores were identified, selected, and analyzed with custom MATLAB scripts.

Hairpin unfolding and refolding events were detected using AutoStepFinder^46^ to identify abrupt changes to the x-position of the anchor fluorophore over time. Extension changes were calculated from the difference in average x-position of the ten frames immediately preceding and following the identified transition. Force exerted on the hairpin at every frame was determined using both length-based and shear stress-based measurements. For length-based force measurements, the total DNA length was calculated at every frame from the Euclidean distance between the localizations of each fluorophore and converted to a corresponding force from the length-to-force relationship determined from our model (Fig. 2l). For shear stress-based measurements, the flow rate in the chamber at every frame was converted to wall shear stresses using the following equation^47^:

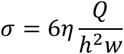

Where *η* is dynamic viscosity, *Q* is volumetric flow rate, *h* is channel height, and *w* is channel width. The determined values of shear stress were then used to calculate corresponding forces from the shear stress-to-force relationship determined by our model (Fig. 2k). Instantaneous loading rates at unfolding and refolding events were determined from the instantaneous rates-of-change of force from both the length and shear stress-based force measurements (with smoothing applied to the length-based force measurements).

FRET traces were extracted from acquired videos using the iSMS software^48^ with donor leakage and γ correction parameters determined from acceptor photobleaching events. For DNA unzipping and Dig-AntiDig rupture data, rupture forces were extracted from acquired videos using a custom MATLAB app. The app automatically identified the frames at which manually selected molecules ruptured from abrupt changes in fluorescence intensity, and correlated the frame to a corresponding force from the known shear stress ramp input into the syringe pump.

### Modelling Flow-Stretched DNA

Our model of flow-stretched DNA builds on theoretical treatments of surface-tethered polymers in shear flow^36,49–51^, and uses empirical parameters fit to experimental data. The DNA contour is discretized into *N* segments of length *n* = 150 bp, approximately equal to the persistence length of DNA (*L_p_* ∼ 50 nm)^35,52^, with any remainder assigned to the segment closest to the anchor.

Each segment *i* (Fig. 2g) experiences a drag force (*f_i_*) proportional to wall shear stress (*σ*), segment length (*n*), and its height above the surface (*h_i_*)^36,49–51^:

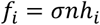

The tension at the beginning of each segment *F_i-1_* is balanced by *f_i_* and the downstream tension *f_i_* (Fig. 2g):

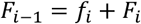

The cumulative tension acting at a segment (*f_i_*) is the sum of drag forces from that segment to the free end (Fig. 2i):

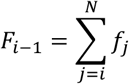

Force at the anchor (*F_0_*) is then obtained by summing all segment drag forces from the anchor to the free end (Fig. 2i):

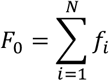

An exact measure of segment height cannot be experimentally determined or calculated from measurable parameters and is thus modelled using a power law function with the parameters *α, β*, and *γ* acquired from fitting to experimental data (Fig. 2j). These parameters describe how height scales with shear stress and the segment’s position along the polymer’s contour *d_i_*, where *d_i_* is the distance along the DNA contour from the free end as a fraction of the total DNA contour length. This flexible form allows independent scaling with shear stress and segment position, allowing for the best fit without artificial coupling of these dependencies.

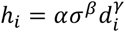

The resulting height profile determined from the optimized model parameters qualitatively agrees with theory from literature on tethered polymers in shear flow, producing a classic “trumpet” profile with a plateau at positions far from the free end, curving upwards near the free end, and gradually flattening with *σ* (Fig. 2h; Fig. S3) ^36,49–51^.

Segment extension (*xi*) is numerically calculated from its tension using the inverse function of the XWLC model. The total expected length (*L*) of the flow-stretched DNA is then the sum of all segment extensions:

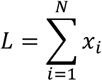

Thus, for any *σ*, the model predicts both *L* and *F_0_*. We determined *α, β*, and *γ* by fitting model-predicted lengths to measured DNA lengths across various shear stresses for two different DNA tethers (8 and 16 kbp, Fig. 2j), using a genetic algorithm that minimized residuals over all calibration datasets. To account for variation in sequence and salt concentrations, an empirical formula for dsDNA persistence length as a function of GC content and ionic strength was used^53^. The globally optimized parameters (*α* = 0.160, *β* = -0.229, and *γ* = -0.157) achieved excellent fits to both DNA tether datasets. The resulting force profiles for both 8 and 16 kbp DNA shows that segment drag forces (*f_i_*) peak at the free end but decrease and plateau toward the anchor (Fig. 2h). This force gradient arises as DNA segments closer to the free end are farther from the surface and thus experience higher flow velocity and greater drag forces (Fig. S3)^36,49^.

### Optical Tweezers Hairpin Construct Preparation

Hairpins used in the optical tweezers experiments were constructed from commercially synthesized DNA hairpins ligated to kilobase-length dsDNA handles. All primer and hairpin sequences are listed in Supplementary Table 1.

The 1019 bp left-handle was generated by PCR amplification of a pBR322 plasmid fragment (New England Biolabs, Whitby, Canada) using Taq DNA polymerase (FroggaBio, Toronto, Canada). The forward primer carried a 5′ biotin modification. The PCR product was digested with PspGI (New England Biolabs) at 75 °C for 1 h to produce a sticky end.

The 1731 bp right-handle was amplified from the Lambda genome (New England Biolabs) with a forward primer carrying a 5′ digoxigenin modification, followed by digestion with TspRI at 65 °C to generate a sticky end.

Both digested handles and the target hairpin were ligated in a one-pot reaction with T4 DNA ligase (New England Biolabs) at room temperature overnight. The resulting DNA construct, consisting of the 5’ biotin-modified left handle, hairpin, and 5’-digoxigenin-modified right-handle, was then gel-purified using a Monarch DNA Gel Extraction Kit (New England Biolabs, Whitby, Canada) and stored at -20°C.

Just prior to the pulling experiment, the DNA construct was incubated with streptavidin-coated 1050 nm diameter polystyrene beads (Bang Labs, Fishers, IN, USA) for at least 30 minutes at room temperature. AntiDig functionalized beads were prepared by incubating 10 μg/mL AntiDig with 1050 nm Protein A-coated polystyrene beads (Bang Labs, Fishers, IN, USA) overnight at room temperature before storage at 4°C. Both types of beads subsequently had their buffers exchanged into the final experimental buffer. All optical trap pulling experiments were performed in 200 mM NaCl, and 10 mM Tris-HCl (pH 8.0), along with 20 mM protocatechuic acid (PCA; Sigma Aldrich, Oakville, Canada) and 50 nM protocatechuate-3,4-dioxygenase (PCD; Sigma Aldrich, Oakville, Canada) as an oxygen scavenger system to prevent DNA damage.

### Optical Tweezers Hairpin Pulling Experiments

Optical tweezers pulling was performed in custom microfluidic chambers on a custom-built dual-trap instrument as described previously^54^. For each experiment, a streptavidin-coated bead incubated with the biotinylated DNA construct and an anti-digoxigenin functionalized bead were trapped independently. Each bead was calibrated individually by collecting position data at 100 kHz for 1 minute and analyzed using custom MATLAB code based on Berg-Sørensen & Flyvbjerg’s optical tweezers power spectrum calibration^55^. The calibrated beads were repeatedly brought into proximity and moved apart until tether formation was observed. The tethers were then pulled by steering the AntiDig-coated bead in a triangle waveform at four different rates in succession – 53, 26.7, 13.3, then 6.7 nm/s – in the same force range until the tether ruptured. OT pulling experiment data on the calibrated beads was collected at 2 kHz at approximately 23-24°C. The 20 bp hairpin dataset was collected over two experimental days; the 15 bp hairpin dataset was collected in a single day.

### Optical Tweezers Analysis

Force-extension curves were processed from raw optical tweezers data as previously described^54^. Multiple tethers were identified and removed manually based on the force-extension signatures. For each valid curve, the contour length at each force was estimated by dividing the measured extension by the XWLC-predicted fractional extension, effectively flattening the curve to facilitate identification of unfolding events. These events were manually reviewed to exclude false positives. The instantaneous loading rate at each unfolding point is calculated by applying a linear fit to the last few points prior to hairpin unfolding in the force-time trace.

