## Supplementary Info for "Massively Parallel Bead-Free Force Spectroscopy with Fluorescence"

### Supplementary Information

| | Page |
|---|---|
| Figure S1 | 2 |
| Figure S2 | 3 |
| Figure S3 | 4 |
| Figure S4 | 5 |
| Figure S5 | 6 |
| Figure S6 | 7 |
| Figure S7 | 8 |
| Figure S8 | 9 |
| Figure S9 | 10 |
| Figure S10 | 11 |
| Table S1 | 12 |
| Supplementary Note 1 | 13 |
| Supplementary Note 2 | 14 |
| Supplementary References | 15 |

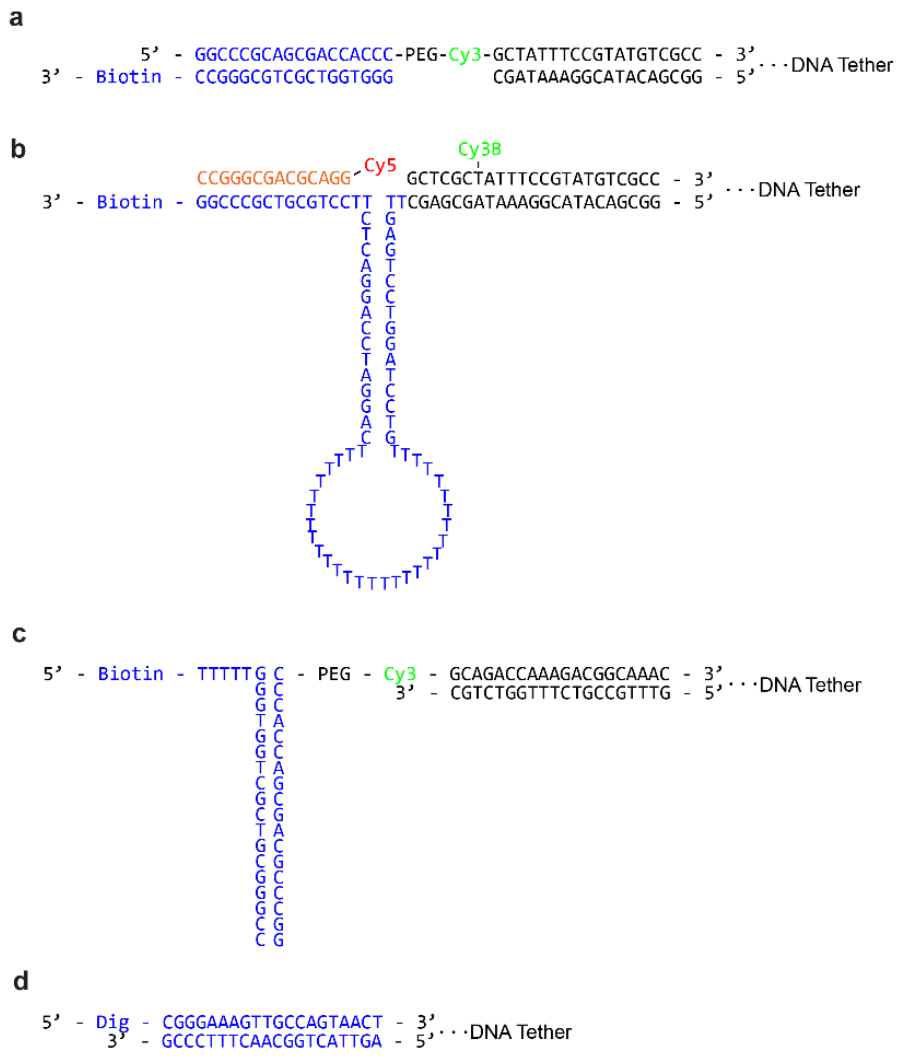


**Figure S1.** Schematic representation of the anchors used in TFS experiments. Calibration experiments performed with both 8 and 16 kbp DNA tethers used a biotinylated 18 bp DNA duplex in the shearing configuration (**a**) as the anchor. The DNA hairpin anchor (15 bp shown) (**b**) was designed to have single stranded sequence complementary to a Cy5-labeled sequence (shown in orange) so that either localization or FRET-based measurements could be obtained. The DNA unzipping anchor (**c**) and the Dig-AntiDig anchor (**d**) were used in the irreversible rupture experiments.

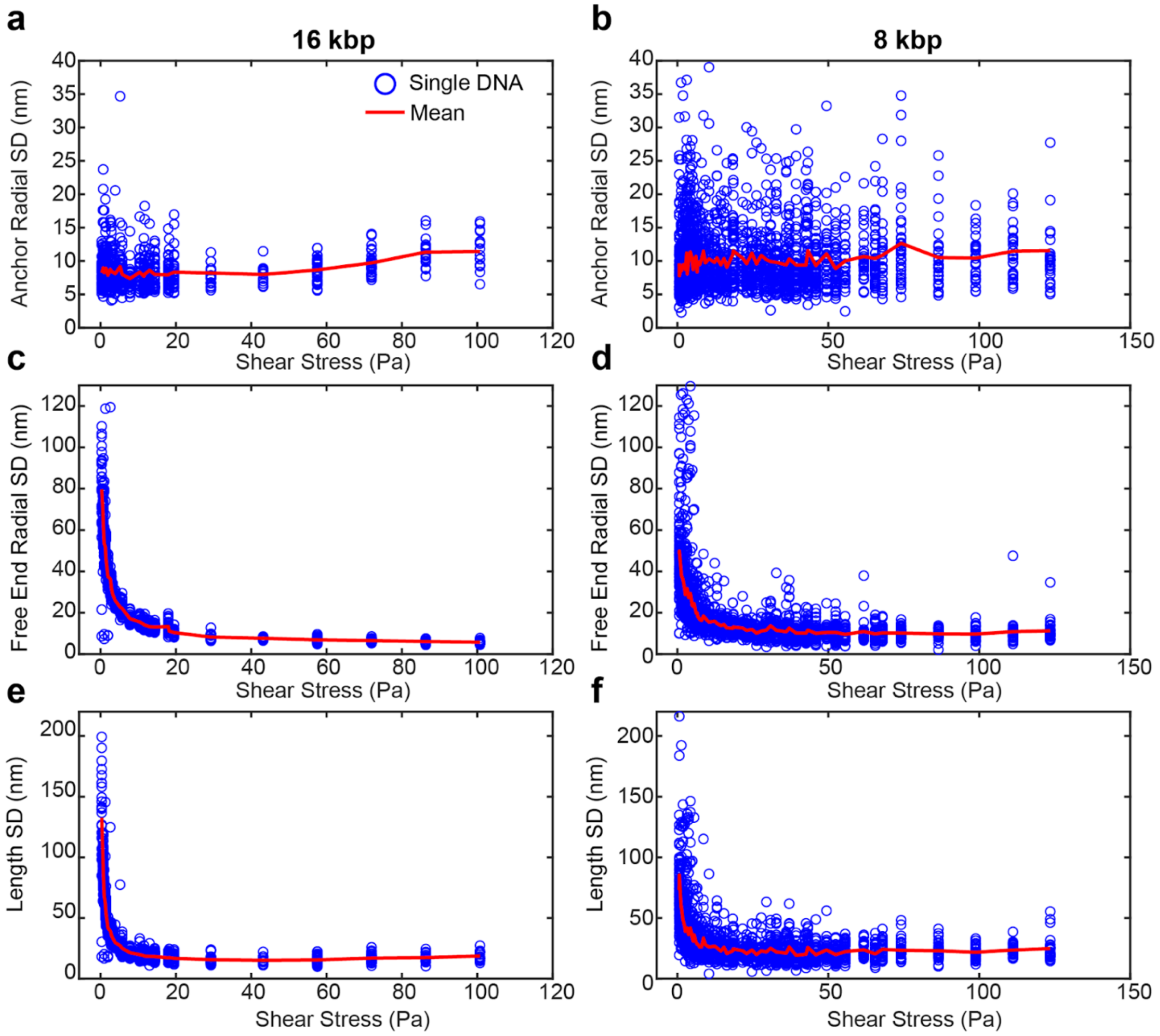


**Figure S2.** Standard deviations of localization positions (**a-d**) and DNA length (**e, f**) as functions of shear stress, based on the calibration data in Fig. 2, for 16 kbp (left) and 8 kbp (right) DNA.

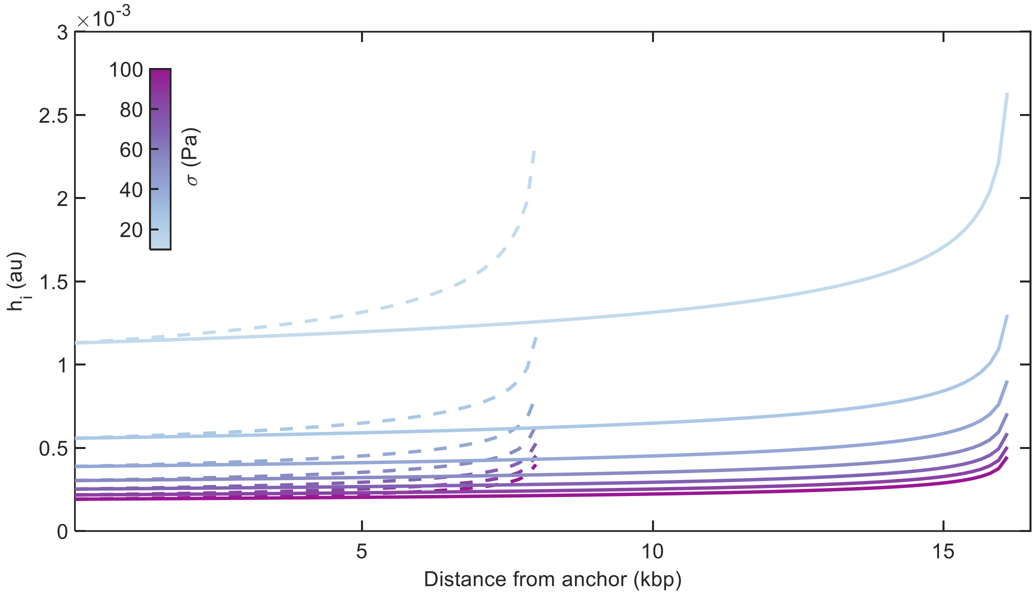


**Figure S3.** Model-predicted height profiles of 8 kbp (dashed) and 16 kbp (solid) DNA across varying shear stresses.

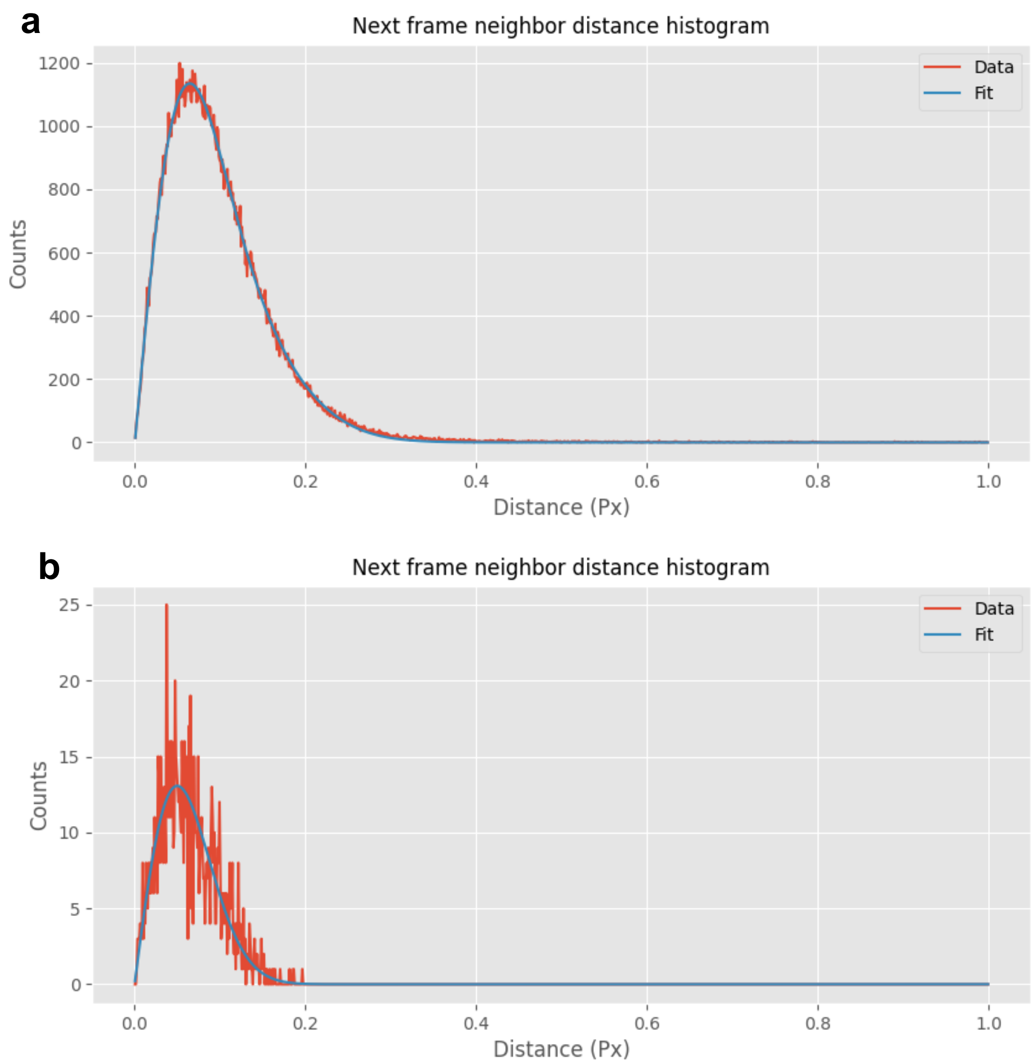


**Figure S4.** Localization precision determined by Nearest Neighbor Analysis (NeNA). (**a**) Nearest-neighbor distance histogram for the entire field of view from the video containing the example trace in Fig. 3b-g. Median fit precision: 3.68 nm; NeNA precision: 8.89 nm. (**b**), Nearest-neighbor distance histogram for anchor localizations in Fig. 3b-g. Median fit precision: 2.36 nm; NeNA precision: 8.02 nm.

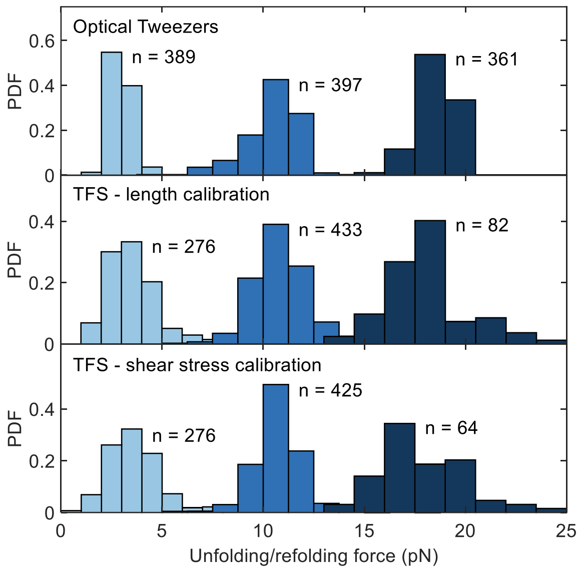


**Figure S5.** Force distributions for 15 bp refolding (left), 15 bp unfolding (center), and 20 bp unfolding (right) measured by optical tweezers (top), TFS localization-based forces (middle), and TFS shear stress–based forces (bottom). Differences in sample size between TFS unfolding and refolding arise from photobleaching events occurring between the two measurements. Differences in sample size between TFS length- and shear stress-based unfolding forces reflect cases where fluorophores photobleached before the force ramp peak was reached; shear stress at each frame was determined by aligning the peak of the stress ramp with the frame of maximum DNA length.

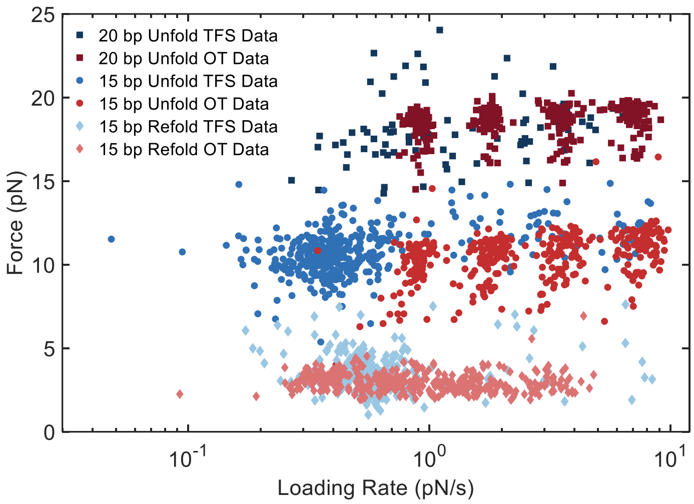


**Figure S6.** Unfolding and refolding forces from optical tweezers and TFS plotted against loading rate on a logarithmic scale, demonstrating result consistencies.

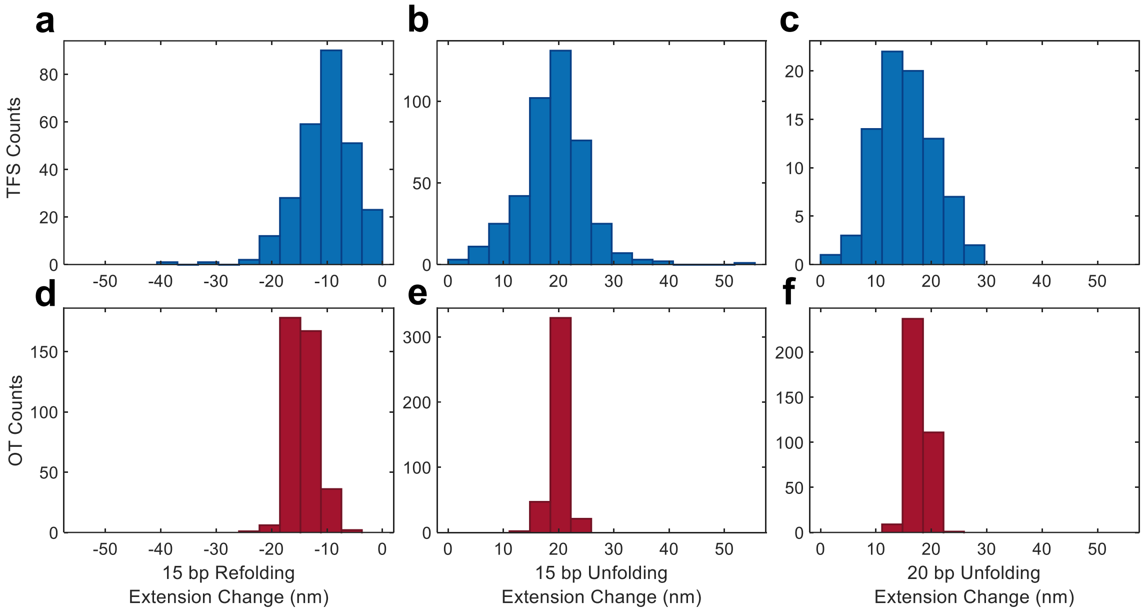


**Figure S7.** Extension change distributions for 15 bp refolding (left), 15 bp unfolding (center), and 20 bp unfolding (right) events measured using TFS (**a-c**) and optical tweezers (**d-f**).

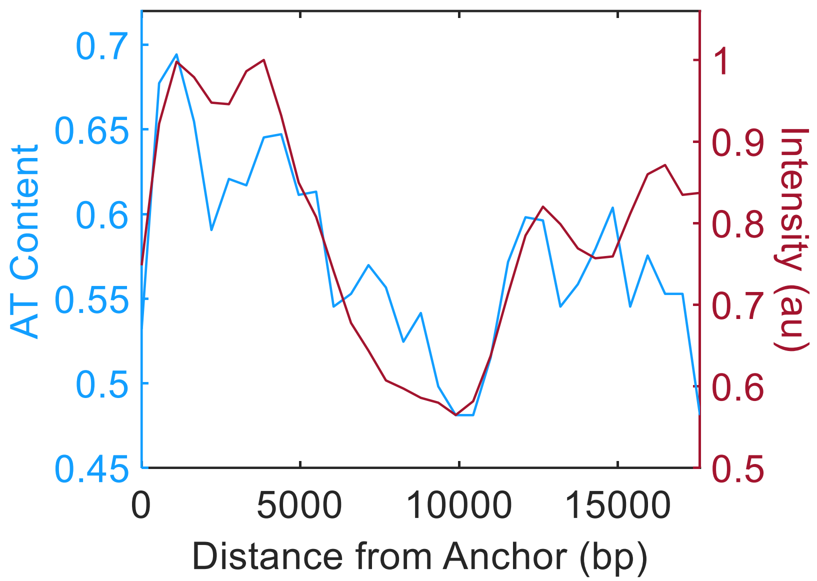


**Figure S8.** AT content overlaid with pixel intensity for the molecule labeled "ii" in Fig. 4c, showing that non-uniform intensity along the DNA is due to preferential SYTOX Deep Red binding to AT-rich regions.

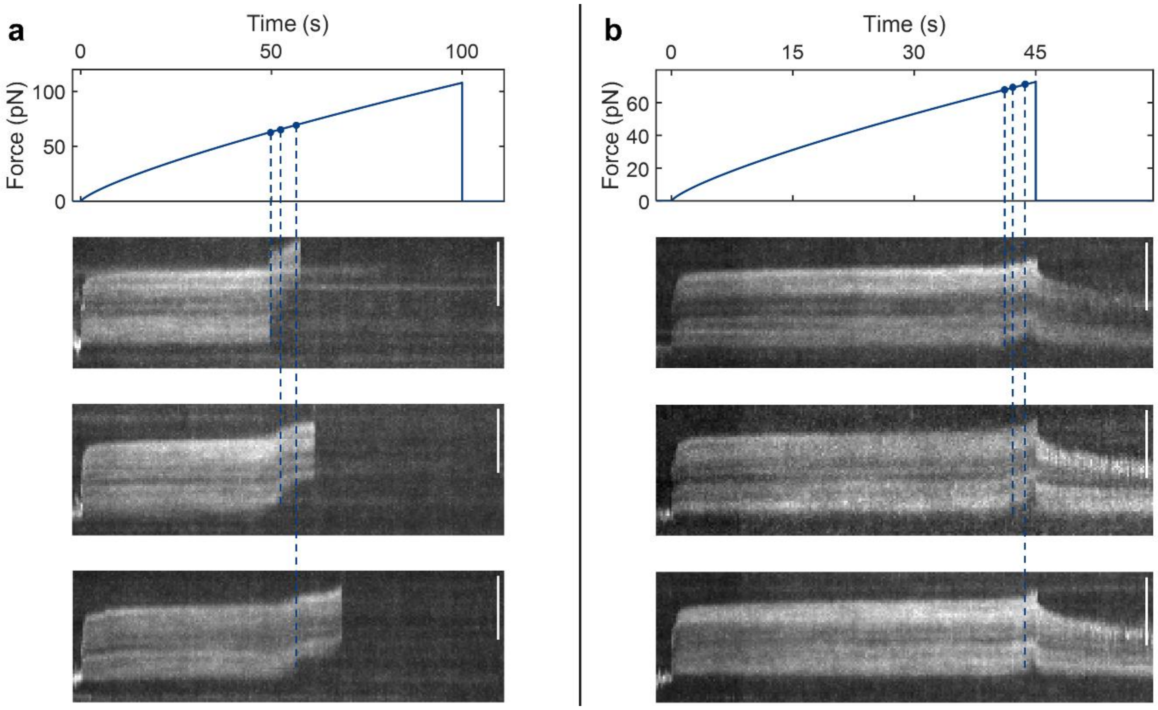


**Figure S9.** Force ramps and corresponding kymographs of 17.6 kbp Dig-modified DNA showing DNA overstretching. Overstretching is identified by rapid fluorescence loss at the anchor end and increased free-end extension, consistent with literature reports for DNA in the presence of intercalating dyes[1]. **a,** Example kymographs (bottom) from a force ramp (top) in which all overstretched molecules subsequently ruptured. **b,** Example kymographs (bottom) from a force ramp (top) in which flow was stopped before Dig-AntiDig rupture, showing molecules retracting to the anchor position in the absence of force, consistent with the reversibility of DNA overstretching. Scale bars are 5 μm.

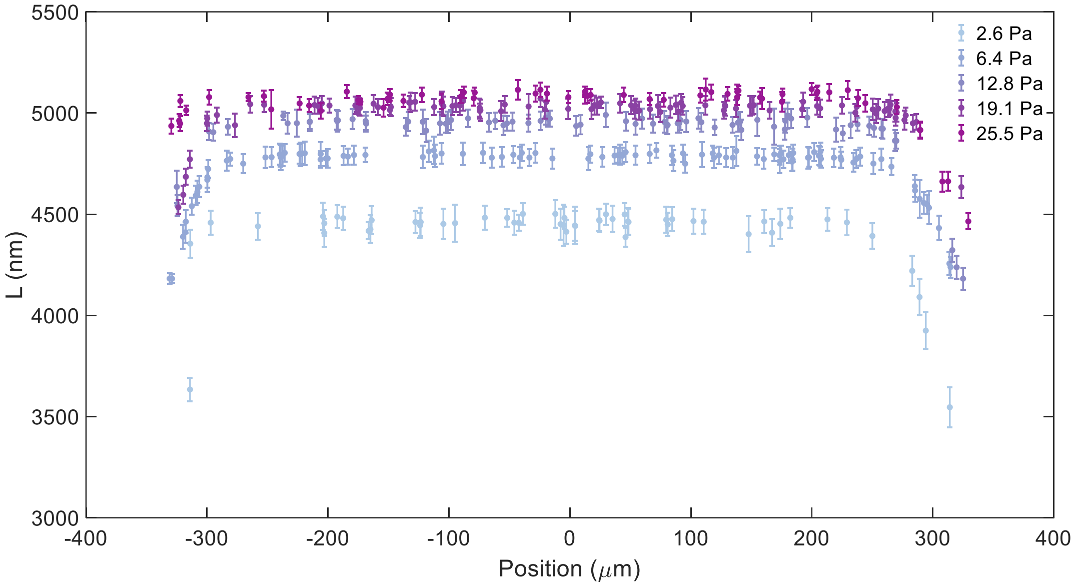


**Figure S10.** Length of 16 kbp DNA at constant shear stresses (2.6-25.5 Pa) plotted against their position across the channel width, showing that ~80% of the central region exhibits consistent stretching. Error bars represent the standard deviation of measured lengths.

**Table S1.** DNA oligonucleotides used in this study. All oligonucleotides were purchased from Integrated DNA Technologies.

| Sequence Name | Sequence (5' → 3') |
|---|---|
| 8 kbp and 16 kbp Calibration Fwd Primer | GGCCCGCAGCGACCACCC /iSp9/ /iCy3/ GCTATTTCCGTATGTCGCC |
| 8 kbp and 16 kbp Hairpin Fwd Primer | GTAGGTTGGTCTCA GCTCGC /iAmMC6T/ ATTTCCGTATGTCGCC |
| 8 kbp Calibration and Hairpin Rev Primer | /5Cy3/ CTTCCGTATCCTTCACCCA |
| 16 kbp Calibration and Hairpin and 12 kbp Unzipping Rev Primer | /5AmMC6/ CCAGGAGGATCTGGAACTTATC |
| 15 bp TFS Hairpin | /5Phos/ GAGC TT GAGTCCTGGATCCTG TTTTTTTTTTTTTTTTTTTTTTTTTTTTTT CAGGATCCAGGACTC TT CCTGCGTCGCCCGG /3Bio/ |
| 20 bp TFS Hairpin | /5Phos /GAGC TT CGCCGCGGGCCGGCGCGCGG TTTTTTTTTT CCGCGCGCCGGCCCGCGGCG TT CCCTCCTGCGTCGCCCGG/3Bio/ |
| 15 bp OT Hairpin | /5Phos/ CCTGG TT GAGTCCTGGATCCTG TTTTTTTTTTTTTTTTTTTTTTTTTTTTTT CAGGATCCAGGACTC TT CCCACTGGC |
| 20 bp OT Hairpin | /5Phos/ CCTGG TT CGCCGCGGGCCGGCGCGCGG TTTTTTTTTT CCGCGCGCCGGCCCGCGGCG TT CCCACTGGC |
| Cy5 Oligo for smFRET 15 bp Hairpin | CCGGGCGACGCAGG /3Cy5Sp/ |
| OT Left Handle Fwd Primer | /5BiosG/ CAGAAGTGGTCCTGCAACT |
| OT Left Handle Rev Primer | CAAGCCTATGCCTACAGCAT |
| OT Right Handle Fwd Primer | /5DigN/ GGGCAAACCAAGACAGCTAA |
| OT Right Handle Rev Primer | CGTTTTCCCGAAAAGCCAGAA |
| 17 kbp Dig Construct Fwd Primer | GTAGGTTGGTCTCAGCTC ATCTCGTTGAAGACCATCGGG |
| 17 kbp Dig Construct Rev Primer | /5DigN/CGGGAAAGTTGCCAGTAACT |
| Surface-Anchoring Unzipping Oligo | /5BiotinTEG/ TTTTT GGGTGGTCGCTGCGGGCC |
| Surface-Anchoring Calibration Oligo | GGGTGGTCGCTGCGGGCC /3BioTEG/ |
| 12 kbp Unzipping Fwd Primer | GGCCCGCAGCGACCACCC /iSp9/ /iCy3/ GCAGACCAAAGACGGCAAAC |

**Abbreviations on oligo modifications (IDT):**
/iSp9/ – triethylene glycol spacer for preserving ssDNA on the 5′ end of a primer
/iCy3/ – internal Cy3
/iAmMC6T/ – internal modified T with a 6C linker to a primary amine for conjugation to Cy3B NHS ester
/5Cy3/ – 5′ Cy3
/5AmMC6/ – 5′ 6C linker to a primary amine for conjugation to Cy3B NHS ester
/5Phos/ – 5′ phosphate modification for ligation
/3Bio/ – 3′ biotin
/3Cy5Sp/ – 3′ Cy5 fluorophore
/5BiosG/ – 5′ biotin
/5DigN/ – 5′ digoxigenin
/5BiotinTEG/ – 5′ biotin linked via triethylene glycol spacer
/3BioTEG/ – 3′ biotin linked via triethylene glycol spacer

**Supplementary Note 1**

The distribution of observed Dig-AntiDig rupture forces contains measurements above DNA overstretching forces (Fig. 4e), as a small fraction of the measured rupture events occurred after overstretching (~15%, see Fig. S9 for example). In the spirit of showing all data and avoiding interpretation bias, these measurements were included in the result (Fig. 4e). However, the reported forces for these events should be interpreted with caution, as partial DNA overstretching can influence force quantification[2]. However, due to the tension gradient along the tether, only the DNA closest to the anchor is overstretched, which is also the DNA that contributes the least to the tension, suggesting that the effect on force quantification may be minor. Regardless, the exact magnitudes of the forces reported in Fig. 4e that are >65 pN should be taken with some degree of uncertainty due to the unknown influence of partial tether overstretching on total force.

Reported Dig-AntiDig dissociation kinetics vary widely in the literature, with zero-force lifetimes ranging from ~67 to ~14,000 s[3–5]. The zero-force lifetime estimated from linear extrapolation in Fig. 4e (~1850 s) falls within this range and is in reasonable agreement with the ~2500 s value predicted from Akbari et al.[5].

**Supplementary Note 2**

The spatial resolution of TFS force–extension measurements depends on the distance measurement method (sm-localization or smFRET) and imaging conditions. For localization-based measurements, we achieved ~8 nm resolution (Fig. S4), while smFRET-based measurements reached ~0.5 nm, as estimated using the method of Holden et al.[6]. These values can be improved by increasing excitation intensity or exposure time to boost photon counts, though this comes at the expense of fluorophore lifetime and temporal resolution, as in conventional single-molecule fluorescence.

The temporal resolution of TFS is ultimately limited by the camera frame rate. Although optical tweezers and AFM can achieve high sampling rate at >10 kHz, TFS's limit is comparable to other video-based SMFS methods and can be increased to 100–1000 Hz using an sCMOS camera with a cropped field of view. Both spatial and temporal resolution can be substantially enhanced using state-of-the-art single-molecule localization methods such as MINFLUX[7], which can achieve <2 nm spatial resolution and 10 kHz temporal resolution with minimal photon flux, enabling extended observation times. Thus, the ultimate spatiotemporal resolution of TFS is dictated entirely by the imaging system.

Force resolution is determined by the precision of either shear stress or DNA length measurements, depending on the calibration used. In a given channel, shear stress accuracy is constrained by syringe pump flow-rate variability (0.05%, <0.05 pN) and channel geometry. Across 80% of the channel width, shear stress is uniform (Fig. S10). Since shear stress scales with $1/h^2$, the ~0.5 μm uncertainty in channel height is the largest contributor to imprecision, corresponding to 1–2% depending on the force range. For length-based force measurements, resolution is limited by fluorophore localization precision, particularly at the free end where Brownian motion contributes additional uncertainty. This effect diminishes as force increases (Fig. S2), improving length-calibration precision from ~15% at ~1 pN to ~5% above 5 pN. The two calibrations are complementary: shear stress calibration offers higher precision but is susceptible to systematic errors from inaccurate channel dimension or flow measurements, whereas length calibration is internally referenced to each molecule's behaviour, avoiding such systematic errors but with lower precision. Using both calibrations in parallel increases confidence in force estimates.

Both force calibrations methods demonstrated sub-pN accuracies within the range of hairpin unfolding and refolding forces. This was confirmed by equivalence testing (two one-sided t-tests $p < 0.05$ for all TFS and OT force distributions in Fig. 3o-q, and Fig. S5, within a threshold of ±1 pN).
